# Polygenic evolution towards a morning chronotype in modern humans

**DOI:** 10.64898/2026.11.01.763544

**Authors:** Shoko Hamamoto, Yuka Moriya, Naoko T. Fujito, Shigekazu Higuchi, Kosuke M. Teshima, Toshiyuki Hayakawa

**Author notes:** Address correspondence to Toshiyuki Hayakawa, Kyushu University, 744 Motooka, Nishi-ku, Fukuoka 819-0395, Japan.

## Abstract

Chronotype is a behavioural phenotype, and its phenotypic trend is considered to change at higher latitudes. In the current study, we performed a polygenic assessment of chronotype trend using associated single nucleotide polymorphisms identified by a genome wide-association study. We found that the phenotypic trend towards morningness increased drastically during the migration of anatomically modern humans above 20 degrees north. Furthermore, two types of latitudinal selection of morningness alleles contributed to establishing the trend towards the morning chronotype at higher latitudes. We propose that the trend towards morningness is stronger in present-day modern humans living above 20 degrees north than in Neanderthals.

## Introduction

Anatomically modern humas (AMHs; *Homo sapiens*) expanded their habitat to almost every part of the planet after emerging in Africa^1^. During this expansion, adaptation to new environments occurred^2^. Chronotype is a behavioural phenotype that is considered to be associated with the latitudinal environment^3^. Previous studies reported a morning to evening latitudinal cline of chronotype towards higher latitudes^4^ and a correlation between evening orientation and higher latitudes^4,5^. These findings suggest an association between the trend towards evening chronotype and higher latitudes^4,5^. In addition, latitude-driven adaptation has been proposed on the basis of an association of allele frequencies of circadian genes and latitude^6^. However, morningness alleles of some genetic loci associated with chronotype were derived from archaic humans through introgression, and their frequencies were found to increase at higher latitudes^7^. Unlike earlier studies, this finding suggests an association between the trend towards a morning chronotype and higher latitudes. Although these previous studies consistently indicate that chronotype is associated with latitude^3-7^, the direction of the reported association differs between studies. In the current study, we hypothesised that a genetic trend towards a morning chronotype at higher latitudes underlies the environmentally induced eveningness orientation. Recently, a genome-wide association study (GWAS) identified 351 genetic loci associated with chronotype (CHR)^8^, making it possible to evaluate its overall polygenic trend. Using these loci, we investigated the adaptive shift in chronotype trend during migration to higher latitudes and examined its evolutionary significance in AMHs by comparing the trend with that of Neanderthals.

## Results

### Chronotype trends and latitudes in AMHs

According to Hu et al.^9^, the effect sizes of associated genetic loci for quantitative phenotypes are well conserved between European ancestry and African ancestry. To assess the phenotypic trend of chronotype in AMHs, we therefore assumed that effect sizes would be highly similar among populations of present-day modern humans (pdMHs). We then applied polygenic score (PGS) and population polygenic score (PPS; an average of PGS values in a population)^10^. We used populations from the 1000 Genomes Project^11^ after removing populations with ambiguous original location (see Materials and Methods).

Fig. 1 shows the distribution of each score for morningness (PGSm and PPSm) along the latitudes of the sample origins (Fig. 1; see Materials and Methods). A significant association between the trend towards a morning chronotype and latitude (α = 0.05/5; *P* < 0.01) was detected in the distributions of PGSm and PPSm (*ρ* = 0.58, *R*^2^ = 0.34, *P* = 0.003) (Fig. 1A and B). This suggests that the trend towards a morning chronotype is associated with migration to higher latitudes. However, several populations from South Asia, Southeast Asia, and the Americas showed high PPSm even though these individuals lived at low latitudes (< 15° N) (Fig. 1B). Considering the route of the global spread of AMHs^1^, such populations underwent a migration to their current habitats at lower latitudes after leaving Africa. If the increase in morningness alleles is adaptive only for migration to higher latitudes and is neutral for migration to lower latitudes, PPSm of such populations would be expected to remain high regardless of adaptation. The exclusion of populations showing migration to lower latitudes mainly revealed a much stronger association between the trend towards a morning chronotype and higher latitudes (*ρ* = 0.90, *R*^2^ = 0.81, *P* = 1.06 × 10^−5^) (Fig. 1C and D). This finding further supports the trend towards a morning chronotype during migration to higher latitudes. Interestingly, when focusing on non-African populations, the association became nonsignificant (*ρ* = 0.75, *R*^2^ = 0.56, *P* = 0.034) (Supplementary Fig. 1). Thus, we concluded that the strong association with latitude depended on a drastic increase in PGSm and PPSm at around 20°N. On the basis of these findings, we focused on the set of populations that did not undergo migration to lower latitudes in our further analyses.

**Fig. 1.**
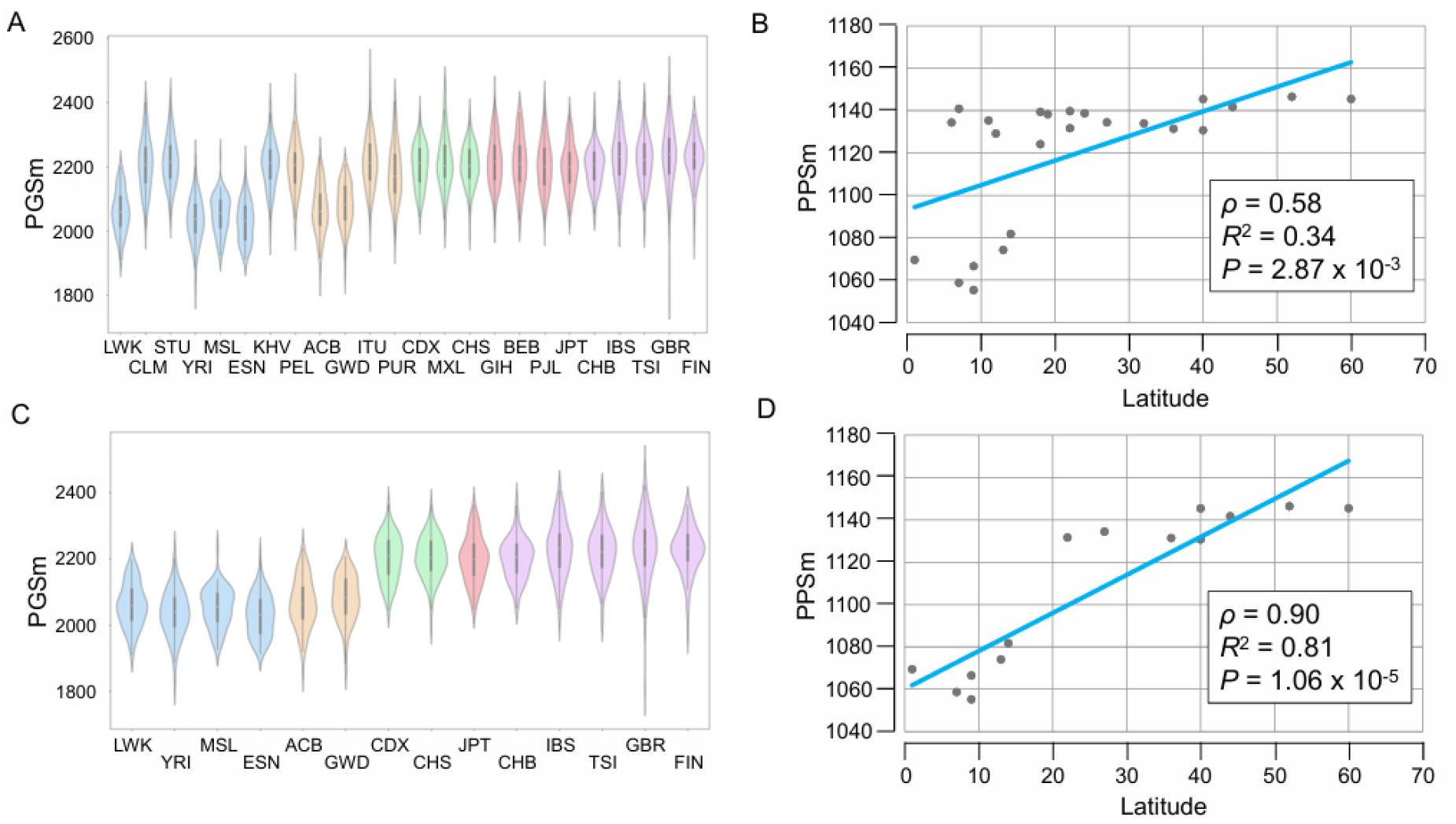
Distribution of polygenic scores along latitudes in AMH populations. (A) PGSm distributions. (B) PPSm distributions. (C) PGSm distribution after the exclusion of populations with migrations into low latitudes. (D) PPSm distribution after the exclusion of populations with migrations into low latitudes. Data are shown for 24 populations from the 1000 Genomes Project^11^: Africa [LWK, ESN, YRI, GWD, MSL, and ACB], Europe [TSI, GBR, IBS, and FIN], South Asia [PJL, GIH, ITU, BEB, and STU], East and Southeast Asia [CDX, KHV, CHS, CHB, and JPT], and the Americas [MXL, CLM, PUR, and PEL].

### Stepwise latitudinal change towards a morning chronotype

To examine the stepwise change towards a morning chronotype with higher latitudes, we divided the populations into five latitudinal groups at 10° intervals: 0–9°, 10°–19°, 20°–29°, 30°–39°, and ≥ 40°. The distribution of PGSm showed differences in the trend towards a morning chronotype among latitudinal groups (Fig. 2). Specifically, the PGSm distribution shifted higher in three groups above 20°N compared with the two groups below 20°N (*P* = 1.86 × 10^−176^; Kruskal-Wallis test for all latitudinal groups; Fig. 2). Furthermore, the trend towards a morning chronotype was further strengthened in the ≥ 40° latitudinal group (*P* = 8.77 × 10^−4^; Kruskal-Wallis test for the 20°–29°, 30°–39°, and ≥ 40° latitudinal groups; Fig. 2). On the basis of the strength of this trend, we propose that the trend towards a morning chronotype increased step-by-step at approximately 20°N and 40°N during migration to higher northern latitudes. Notably, even when populations that migrated to lower latitudes were included, a similar trend towards a morning chronotype was suggested (see Supplementary Fig. 2).

**Fig. 2.**
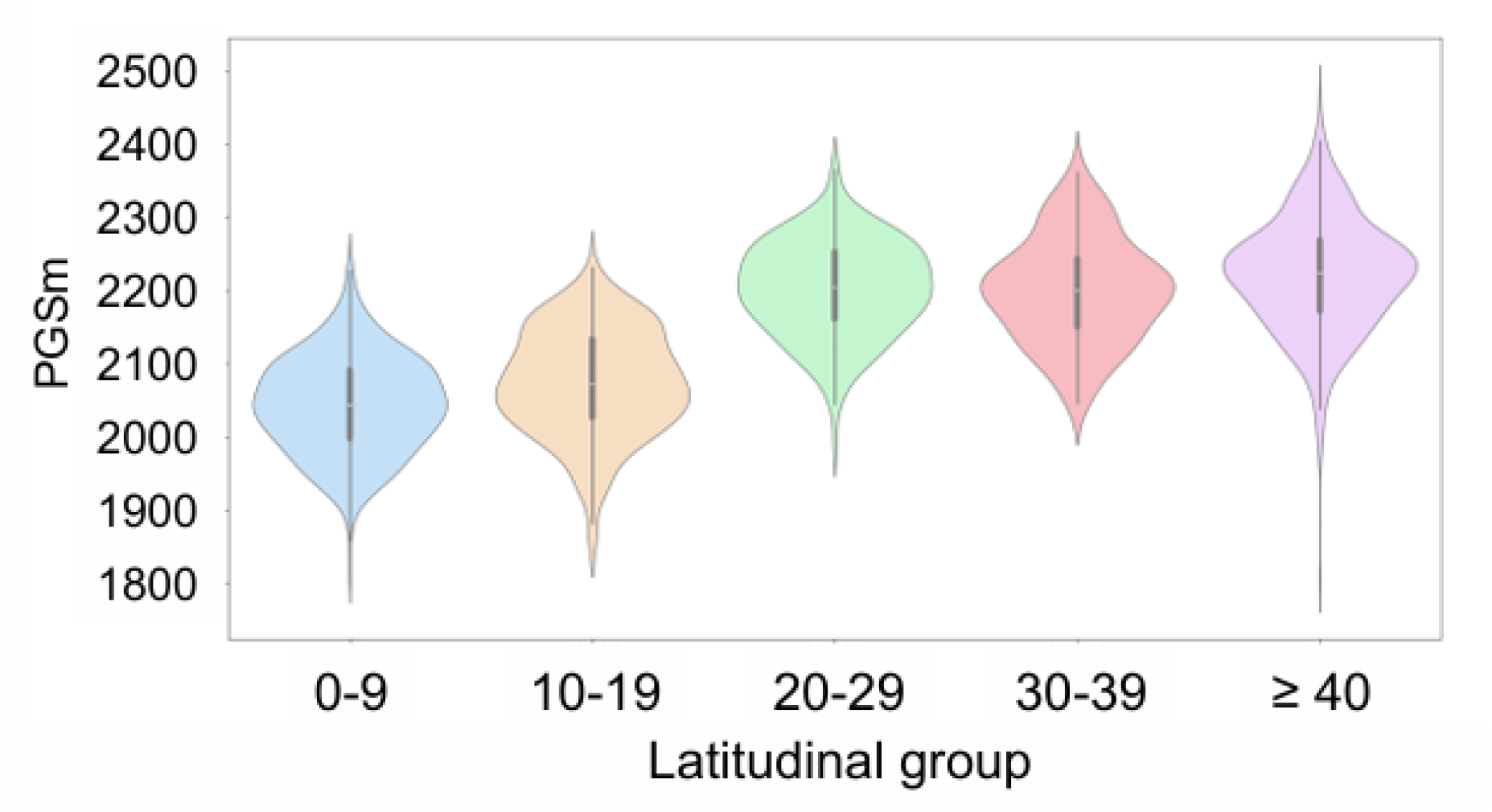
Distributions of PGSm in latitudinal groups. Populations belonging to each latitudinal group are as follows: LWK, YRI, MSL, and ESN for 0°–10°; ACB and GWD for 10°–19°; CDX and CHS for 20°–29°; JPT for 30°–39°; CHB, IBS, TSI, GBR, and FIN for ≥ 40°. The Bonferroni-corrected threshold is *P* = 0.0125 for four Kruskal-Wallis tests (see also Supplementary Fig. 2).

To examine whether the observed stepwise increase in PPSm, particularly around 20°N, is a general feature among associated loci identified by GWAS, we also analysed height-associated SNPs (HEI)^12^, because HEI was obtained by one of the largest GWASs^12^. HEI showed no stepwise change in the PPS of positive-effect alleles (PPSp) at around 20°N (Supplementary Fig. 3A). Thus, the results suggested that the stepwise change towards morningness is characteristic of CHR. Interestingly, HEI indicated that populations from Europe have a high PPSp (Supplementary Fig. 3A). Height is significantly influenced by environmental factors such as nutrition. By focusing on PPSp values from developed countries, we found that PPSp values in populations from Europe were much higher than those from East Asia (Supplementary Fig. 3B). Because this is consistent with the well-known trend in height (i.e., greater height in Europe than in East Asia)^13^, this supports the applicability of PPS for assessing relative differences in phenotypic trends at the population level, provided that the extent of such differences is primarily determined by genetic rather than environmental differences.

### Increase of morningness alleles during migration to above 20°N

In particular, the higher PPSm in populations living above 20°N compared with those living below 20°N raised the possibility of adaptive evolution towards a morning chronotype at those latitudes during AMH migration. To test this possibility, we compared the proportion of alleles showing higher frequencies in populations living above 20°N than in those living below 20°N between morningness and eveningness alleles. Consequently, this proportion was significantly larger for morningness alleles (140/184 in derived alleles; 49/148 in ancestral alleles) than for eveningness alleles (99/148 in derived alleles; 44/184 in ancestral alleles) (*P* = 0.004 for derived alleles; *P* = 0.007 for ancestral alleles; binomial test; Bonferroni-corrected threshold of *P* = 0.025) (Supplementary Fig. 4). Because lower allele frequencies in populations living below 20°N were associated with higher frequencies in those living above 20°N (*ρ* = 0.55, *R*^2^ = 0.30, *P* = 5.79 × 10^−16^ for the derived morningness alleles; *ρ* = 0.54, *R*^2^ = 0.29, *P* = 1.55 × 10^−12^ for the derived eveningness alleles), it is possible that this difference in rate might be caused by a higher proportion of low-frequency alleles among morningness alleles compared with eveningness alleles in populations living below 20°N. However, no such difference in the frequency distribution within CHR was detected in populations living below 20°N (*P* > 0.05; Supplementary Fig. 5). Thus, the greater increase of frequencies in morningness alleles than in eveningness alleles at around 20°N suggests that a selective increase in morningness alleles occurred at around 20°N during the spread of AMHs. Taken together with the drastic increase in PPSm, these findings demonstrate adaptive evolution towards a morning chronotype at around 20°N.

### Selective increase of morningness alleles along higher latitudes

If a certain allele is favoured by adaptation to higher latitudes, its frequency would be expected to increase at higher latitudes. Of the 332 associated loci, 62 morningness and 46 eveningness alleles showed a continuous frequency increase along higher latitudes (i.e., from below 20°N to 20–39°N, and above 40°N). This significantly greater number of morningness alleles (*P* = 0.009; binomial test) suggests the involvement of a selective increase in morningness alleles with migration to above 40°N. To examine this possibility, we performed *F*c analysis to detect signals of positive selection in individuals from Europe, whose populations examined in this study originated from locations at or above 40°N^14–16^. Among the 62 morningness alleles showing a continuous increase along higher latitudes (hereafter referred to as latitudinal morningness alleles), we identified signals of positive selection on two alleles at rs4027217 (*F*_c_ = 0.070, *P* = 1.0 × 10^−4^) and rs6727752 (*F*_c_ = 0.000, *P* = 1.0 × 10^−4^). This finding raises the possibility that the selective increase of specific morningness alleles contributed to the highest PPSm in the population at or above 40°N. Interestingly, the distributions of morningness allele frequencies differed between rs4027217 and rs6727752 (Fig. 3). The former distribution showed a drastic increase at around 20°N (Fig. 3A), similar to the PPS distribution (Fig. 1D). In contrast, the latter showed a relatively linear increase of morningness alleles along higher latitudes (Fig. 3B). This finding suggests that these two types of latitudinal increase in morningness alleles underlie the adaptive evolution shaping the PPSm distribution.

**Fig. 3.**
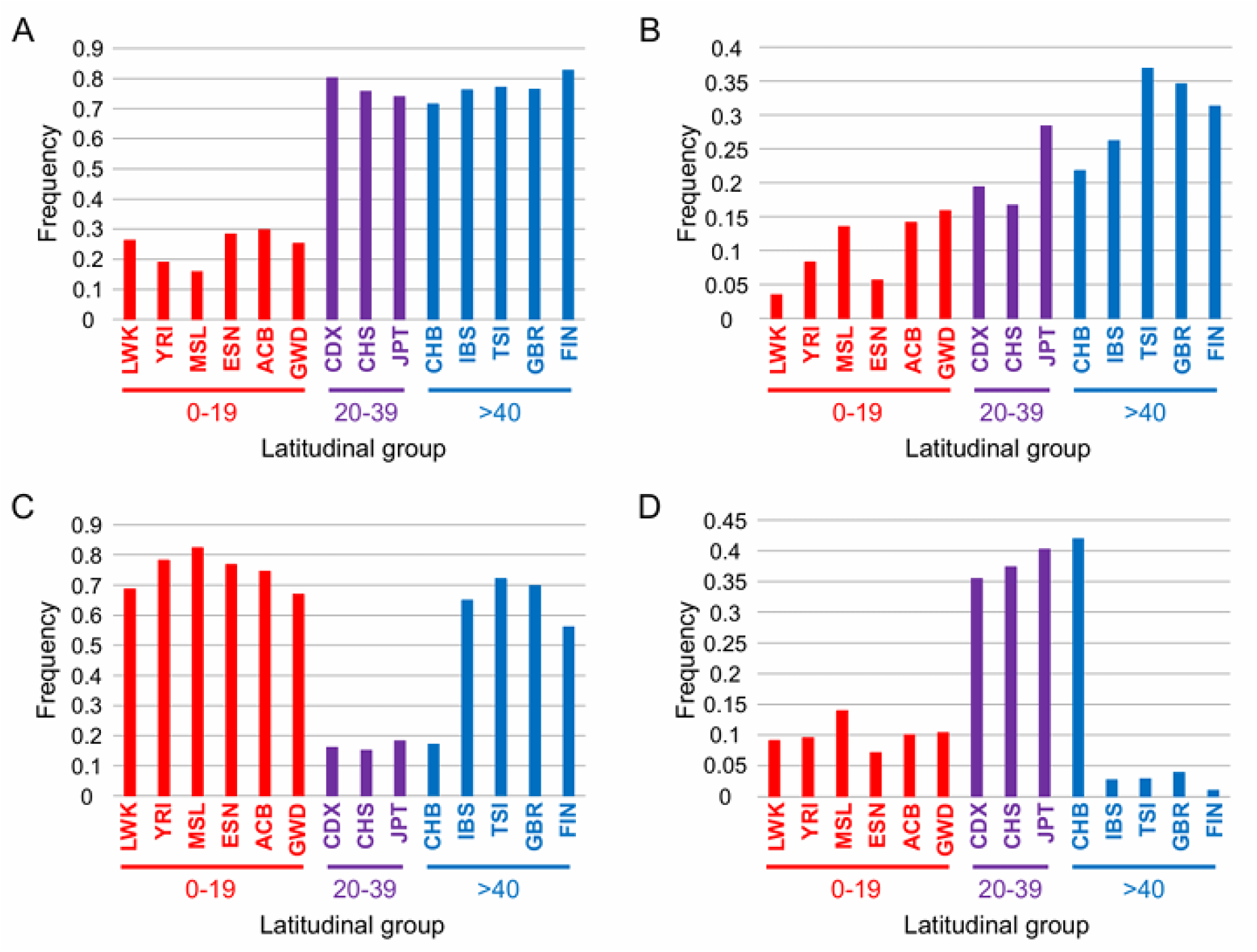
Distribution of allele frequencies. (A) Morningness allele of rs4027217. (B) Morningness allele of rs6727752. (C) Morningness allele of rs17712705. (D) Eveningness allele of rs5016898. Data are shown for populations from the 1000 Genomes Project^11^.

### Latitudinal selection of morningness alleles

Interestingly, the PPSm distribution was similar to that obtained with homogeneous effect sizes (Supplementary Fig. 6). This suggests that the PPSm distribution mainly depends on the frequencies of morningness alleles. To elucidate how the selective increase of morningness alleles, exemplified by rs4027217 and rs6727752, was associated with their overall evolution, we used the frequency distributions of these two alleles as yardsticks. We then examined the correlation in the morningness alleles between correlation coefficient (*ρ*) values with the yardsticks and the average *ρ*-values with other morningness alleles. Bonferroni-corrected significant positive correlations (α = 0.05/6; *P* < 0.008) were found in every yardstick (Fig. 4A and B). Higher *ρ*-values with the yardstick were associated with higher average *ρ*-values with other morningness alleles in the morningness alleles (*R*^2^ = 0.94, *P* = 7.2 × 10^−203^; slope = 0.10 for rs4027217; and *R*^2^ = 0.98, *P* = 4.9 × 10^−279^; slope = 0.11 for rs6727752) (Fig. 4A and B). A similar significant association was found in the latitudinal morningness alleles (*R*^2^ = 0.30, *P* = 5.9 × 10^−6^; slope = 0.34 for rs4027217; and *R*^2^ = 0.62, *P* = 7.4 × 10^−14^; slope = 0.77 for rs6727752) (Fig. 4C and D). These results indicate that a trend towards becoming similar to the yardsticks was involved, especially in the evolution of the latitudinal morningness alleles. This suggests that two types of latitudinal selection of morningness alleles contributed to the trend towards a morning chronotype at higher latitudes. Unfortunately, we could not elucidate the biological functions behind these two latitudinal selections because rs4027217 and rs6727752 are not located on known genes.

**Fig. 4.**
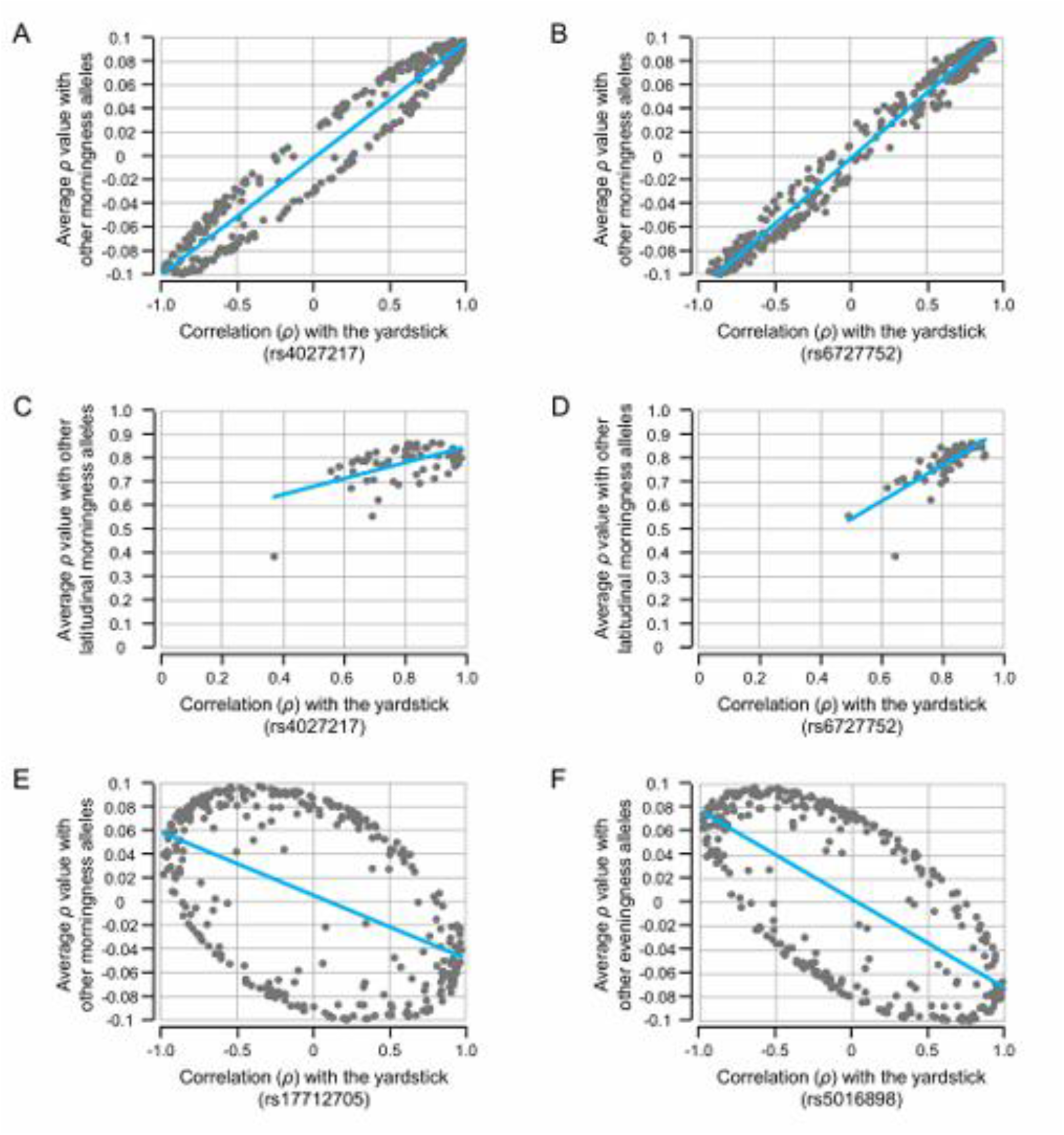
Association with the yardsticks. (A) Association with the morningness allele of rs4027217 in overall morningness alleles. (B) Association with the morningness allele of rs6727752 in overall morningness alleles. (C) Association with the morningness allele of rs4027217 in the latitudinal morningness alleles. (B) Association with the morningness allele of rs6727752 in the latitudinal morningness alleles.

### Inclusion of alleles negatively associated with chronotype evolution

In addition to rs4027217 and rs6727752, signals of positive selection were detected for the morningness allele at rs17712705 (*F*_c_ = 0.045, *P* = 1.0 × 10^−4^) and the eveningness allele at rs5016898 (*F*_c_ = 0.006, *P* = 1.0 × 10^−4^). We examined the degree of the association of selective allele increase shown by rs17712705 and rs5016898 with the evolution of chronotype-associated alleles. Unlike with rs4027217 and rs6727752, the frequency distribution of the morningness allele at rs17712705 was negatively associated with those of other SNPs (*R*^2^ = 0.23, *P* = 1.8 × 10^−20^; slope = −0.05) (Fig. 4E). Meanwhile, the frequency distribution of the eveningness allele at rs5016898 was also negatively associated with those of other SNPs (*R*^2^ = 0.43, *P* = 3.6 × 10^−42^; slope = −0.07) (Fig. 4F). The observed negative associations suggest that the selective increases in these alleles are not associated with the trend towards a morning chronotype at higher latitudes.

### Comparison of chronotype trend between AMHs and Neanderthals

To examine polygenic differences in chronotype trends between AMHs and Neanderthals, we next compared their PPSm values. All genetic loci associated with chronotype were obtained in the Neanderthal individuals examined in our study. In the present study, we examined five Neanderthal individuals whose whole-genome sequences are of high quality and high coverage. These individuals were as differentiated as the most differentiated present-day human populations worldwide^17^. The PGSs of the five individuals were very similar to each other, and their divergences were much smaller than those in pdMHs from Europe (Supplementary Fig. 6). The effect sizes of genetic loci associated with chronotype showed a distribution biased towards small values (Supplementary Fig. 7), indicating that phenotype expression depends on the orchestration of multiple genetic loci with small effect sizes. Even if Neanderthals had unique effect sizes at each genetic locus, any differences with those of AMHs would be expected to be negligible.

Before comparing PPSm, we tested whether the PPSm values estimated from five individuals could reproduce the actual PPSm values of a population. First, we generated a distribution of estimated PPSm by randomly selecting five individuals from Europe 10,000 times and compared this distribution with the actual PPSm of Europe. Although the mode of the generated distribution matched well with the actual PPSm, the reproduction accuracy of the actual PPSm was low (∼7%) (Supplementary Fig. 8), indicating that PPSm from five individuals is not an appropriate proxy for actual PPSm. Therefore, we next selected a number of alleles from Europe for each SNP that was equal to the number of alleles available from five Neanderthal individuals, and used a possible distribution inferred from the frequencies of morningness alleles in the selected alleles (Materials and Methods). Even when the comparison between an inferred possible PPSm distribution and the actual PPSm was replicated 10,000 times, the actual PPSm never fell outside the inferred distribution (Supplementary Fig. 9). This indicates that, despite the small sample size of five individuals, evaluating the distribution was equivalent to addressing the actual PPSm. Consequently, for Neanderthal PPSm, we decided to use a possible distribution inferred from the frequencies of morningness alleles in the five Neanderthal individuals (Materials and Methods).

On the basis of the considerations described above, we assumed that the gene repertories and effect sizes were similar between AMHs and Neanderthals, then compared the genetic trends of chronotype between the two human groups using PPSm. When PPSm was estimated using all samples of pdMHs, no significant difference was detected between AMHs and Neanderthals (Fig. 5A). However, when PPSm was calculated separately by latitudinal groups, the 20°–29°, 30°–39°, and ≥ 40° latitudinal groups showed Bonferroni-corrected significance (α = 0.05/22; *P* = 2.3 × 10^−3^), and their PPSm values were higher than those of Neanderthals (Fig. 5B). Even when PPS was estimated with homogeneous effect sizes (Fig. 5C and D), significance was observed in the population above 20°N (Fig. 5D). It is noteworthy that the impact of population structure appeared to be negligible in the PPSm estimation for Neanderthals because of the consistency of the phenotypic trend observed with PGSm (Fig. 5 and Supplementary Fig. 6). Notably, even when populations that migrated to lower latitudes were included, the results remained unchanged (see Supplementary Fig. 10).

**Fig. 5.**
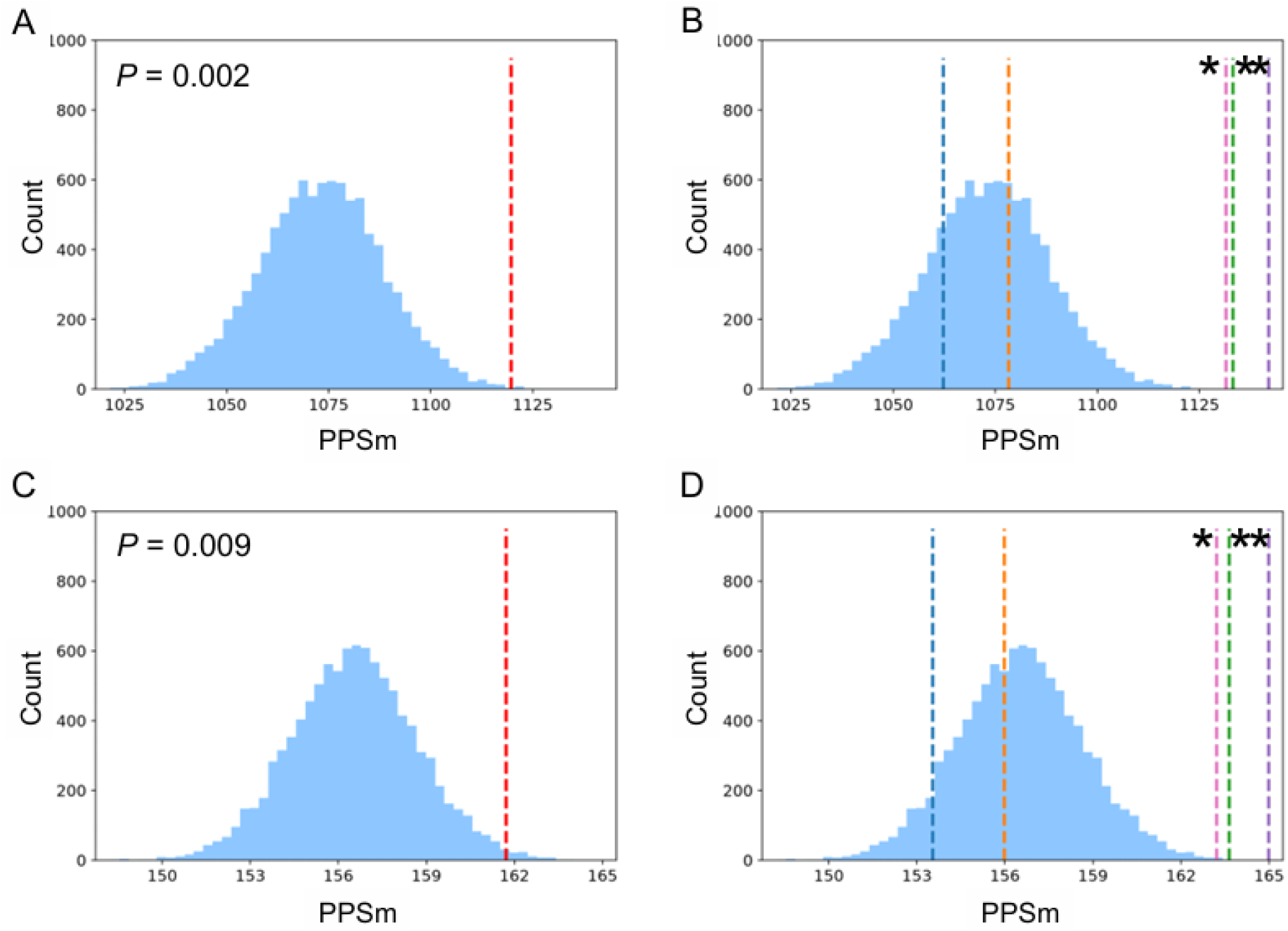
Comparison of chronotype trends between AMHs and Neanderthals. (A, C) PPSm values. PPSm of pdMHs was estimated by excluding populations with migrations into low latitudes and is shown by the red vertical dotted line. The possible distribution of PPSm for Neanderthals is shown in blue. (B, D) PPSm values of the latitudinal groups (0°–9°, 10°–19°, 20°–29°, 30°–39°, and ≥ 40°) were estimated by excluding populations with migrations into low latitudes, and are shown as vertical dotted lines in blue, orange, green, pink, and purple, respectively. The PPSm for Neanderthals is shown by a possible distribution. PPSm estimation was performed with effect sizes from GWAS (A, B) and with homogeneous effect sizes (C, D). Asterisks indicate a Bonferroni-corrected significant difference compared with the Neanderthal distribution.

## Discussion

A drastic increase in PPSm and a selective increase of morningness alleles were detected at around 20°N during the spread of AMHs (Figs. 1 and 2). Because 20°N corresponds approximately to the Tropic of Cancer, the observed adaptive shift towards a morning chronotype can be interpreted as an adaptation to seasonal variations in daylight hours as these variations become more pronounced around the Tropic of Cancer. Unfortunately, it is not clear why the trend towards a morning chronotype would be beneficial at higher latitudes. However, it has been previously reported that morning light exposure has an effect on resetting the circadian clock^18^. Taking this finding into account, a morning chronotype may be helpful to reset the circadian clock by catching morning light in changing daylight hours through the year, and thereby maintain performance in the daytime.

The prevalence of seasonal affective disorder (SAD) has been reported to increase with latitude, and seasonal changes in the light environment, including shorter daylight duration during winter, are considered to contribute to its development^19^. In addition, evening chronotype and delayed sleep phase have been associated with greater seasonality and SAD^20,21^. According to the phase-shift hypothesis of SAD, a delayed circadian phase relative to the sleep–wake cycle may contribute to winter depression, and correction of this circadian misalignment by phase advance has been associated with improvement of depressive symptoms^22^. These findings raise the possibility that, at higher latitudes where seasonal variation in daylight is greater and morning light is limited during winter, a circadian phenotype less susceptible to phase delay, such as morningness, may have been advantageous. Thus, the increase in morningness-associated alleles observed at higher latitudes in the present study may partly reflect adaptation to such seasonal light environments. However, this interpretation remains speculative, and the physiological mechanisms linking morningness to adaptation to high-latitude environments require further investigation.

In contrast to previous studies that reported a trend towards an eveningness chronotype at higher latitudes^4,5^, the current findings support a trend towards a morningness chronotype at higher latitudes. This is consistent with a previous genetic observation of a positive association between higher morningness allele frequencies and higher latitudes^7^. Because chronotype is a behavioural phenotype influenced by societal pressures^8^, the discrepancy among previous reports might indicate that the trend towards a morningness chronotype is an evolutionarily established genetic trend, whereas the trend towards eveningness is driven by recent societal factors (i.e., artificial light exposure at night in urban areas and the susceptibility of different climate zones to urbanisation). Thus, an individual’s chronotype in the present day might be determined primarily by societal factors rather than their underlying genetic predisposition.

Regardless of the effect sizes, the PPSm was significantly higher in pdMHs living above 20°N than in Neanderthals (Fig. 5B and D). This raises the possibility that certain adaptations associated with the trend towards a morning chronotype (e.g., an adaptation to seasonal variations in daylight hours) differed between them, and that Neanderthals were thereby disadvantaged compared with AMHs in living above 20°N.

In conclusion, the findings of the current study suggest that the trend towards a morning chronotype, to which two types of latitudinal selection of morningness alleles contributed, was drastically increased at around 20°N during the AMH migration. These results may suggest that adaptive evolution towards a morning chronotype occurred around the Tropic of Cancer as an adaptation to the seasonal variations of daylight in AMHs. This study also raises the possibility that such a trend differed between AMHs living above 20°N and Neanderthals. This study merely focused on an evolutionarily established genetic trend. Because chronotype is a behavioural phenotype influenced by societal environments, our findings may be helpful for examining the degree of influences of societal factors in an individual’s chronotype by providing an individual’s genetic baseline.

## Materials and Methods

### SNPs associated with chronotype

SNPs associated with chronotype were obtained from GWASs^8^. On the basis of the 1000 Genomes Project database^11^, we eliminated SNPs for which allelic frequencies are not available, SNPs for which multiple derived alleles are represented, SNPs for which only derived alleles are found in AMHs, SNPs located on the X chromosome, and indels.

### DNA sequence data

DNA sequences were retrieved from phase 3 of the 1000 Genomes Project database^11^, covering 2,504 individuals (5,008 sequences) from 26 different global populations.

DNA sequences of five Neanderthals (Vindija 33.19, Denisova 5, Chagyrskaya 8, Denisova 17, and Goyet 1) were obtained from public databases (http://cdna.eva.mpg.de/neandertal/Vindija/; http://cdna.eva.mpg.de/neandertal/altai/; http://ftp.eva.mpg.de/neandertal/Chagyrskaya/VCF/; ftp://ftp.sra.ebi.ac.uk/vol1/run/ERR167/ERR16782339/A26088_all.uniq.L35MQ25.realign.bam; and https://edmond.mpg.de/dataset.xhtml?persistentid=doi:10.17617/3.f9n73o).

### Latitudes

We used the approximate latitude of each population’s sampling location. However, if the sampling site differed from the birthplace or ancestral origin, we used the approximate latitude of the capital city of the country of ancestral origin or the central city of the birthplace. The latitudes used were as follows: 7° for YRI (Yoruba in Ibadan, Nigeria), 1° for LWK (Luhya in Webuye, Kenya), 14° for GWD (Gambian in Western divisions in the Gambia), 9° for MSL (Mende in Sierra Leone), 9° for ESN (Esan in Nigeria), 13° for ACB (African Caribbeans in Barbados), 44° for TSI (Toscani in Italy), 60° for FIN (Finnish in Finland), 52° for GBR (British in England and Scotland), 40° for IBS (Iberian population in Spain), 22° for GIH (Gujarati Indian from Houston, Texas, USA), 32° for PJL (Punjabi from Lahore, Pakistan), 24° for BEB (Bengali from Bangladesh), 7° for STU (Sri Lankan Tamil from the UK), 18° for ITU (Indian Telugu from the UK), 40° for CHB (Han Chinese in Beijing, China), 36° for JPT (Japanese in Tokyo, Japan), 27° for CHS (Southern Han Chinese, China), 22° for CDX (Chinese Dai in Xishuangbanna, China), 11° for KHV (Kinh in Ho Chi Minh City, Vietnam), 19° for MXL (Mexican ancestry in Los Angeles, California), 18° for PUR (Puerto Rican in Puerto Rico), 6° for CLM (Colombian in Medellin, Colombia), and 12° for PEL (Peruvian in Lima, Peru). We eliminated ASW (Americans of African ancestry in Southwest USA) and CEU (Utah residents with Northern and Western European ancestry) because of ambiguity regarding the original locations.

### Population polygenic score (PPS)

PGS is generally applied to assess the degree of phenotypic trend in an individual^23^. On the basis of the PGS, we developed the PPS to assess phenotypic trends at the population level.

PPS is obtained using the following formula:

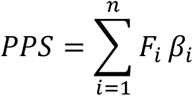

where *n* is total number of associated SNPs, *F* is the frequency of allele at an associated SNP in a certain population, and *β* is the effect size estimated in a GWAS^10^.

### Estimation of PPS as a possible distribution in Neanderthals

Using the beta distribution on the basis of allele frequencies in five Neanderthal individuals, we first obtained a possible distribution of the allele frequency at each genetic locus and subsequently calculated the PPS by randomly obtaining frequency (*F*) from this distribution. We then estimated PPS as a distribution from 10,000 replications of the calculation. All SNPs were available in at least one of the Neanderthal individuals we examined.

### Increase of allele frequencies at 20°N latitude

To compare morningness and eveningness alleles, we calculated the observed number of alleles showing an allele frequency increase at above 20°N latitude. This excluded alleles with a frequency of 0 at < 20°N latitude. The Bonferroni-corrected significance threshold was set to *P* = 0.0125 for derived and ancestral alleles across two GWAS datasets (CHR and HEI).

### Detecting positive selection

We applied the *F*_c_ statistic to detect positive selection on SNPs in three meta-populations: Africa, Europe, and East and Southeast Asia^14–16^. To detect all possible genetic loci, we modified a previously reported simulation method^14,16^ by including alleles showing an *S*- value (number of segregating sites) of at least 100. Of the 332 associated loci, 196 SNPs met the criteria for the *F*_c_ analysis.

## Acknowledgements

This research was supported by JSPS KAKENHI (grant numbers JP16K07535, JP19K06866, and JP24K09628 to T.H.). We thank Benjamin Knight, MSc., from Edanz (https://jp.edanz.com/ac) for editing a draft of this manuscript.

## Author contributions

T.H. conceived and supervised the study. S. Hamamoto, Y.M., N.T.F., and T.H. collected sequence data. S. Hamamoto, Y.M., and T.H. performed analyses of phenotypic trends. N.T.F. and S. Hamamoto performed analysis detecting signals of positive selection. S. Hamamoto, Y.M., N.T.F., and K.M.T. read and commented on the pre-final version of the manuscript.

T.H. and S. Higuchi wrote the paper.

## Data availability

The genomes used are available from the 1000 Genomes Project (phase 3 release, https://www.internationalgenome.org/).

