## Supplementary information for "Polygenic evolution towards a morning chronotype in modern humans"

Supplementary information includes:
Supplementary Figures 1 to 10

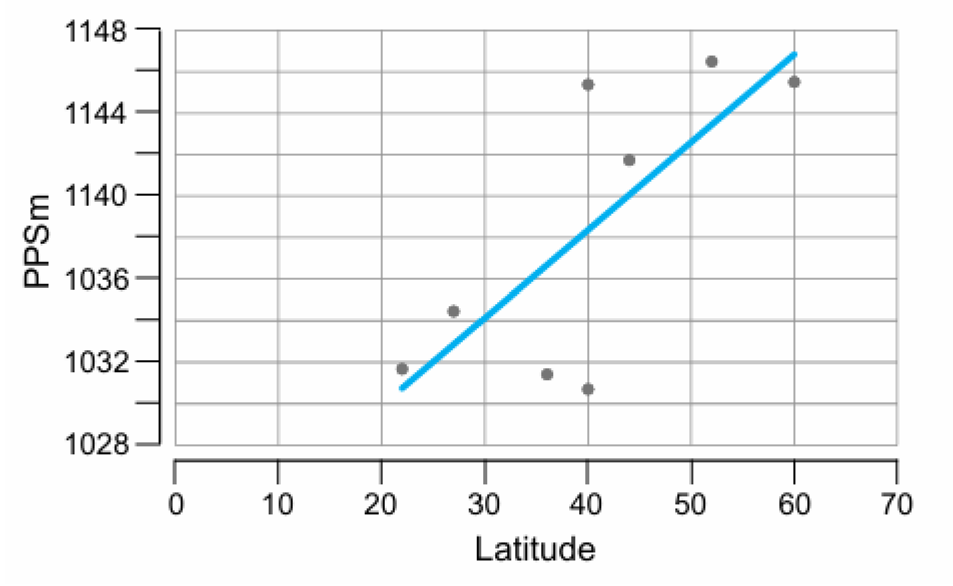


Supplementary Fig. 1. Association between latitude and morningness allele frequencies in non-African populations (CDX, CHS, JPT, CHB, IBS, TSI, GBR, and FIN).

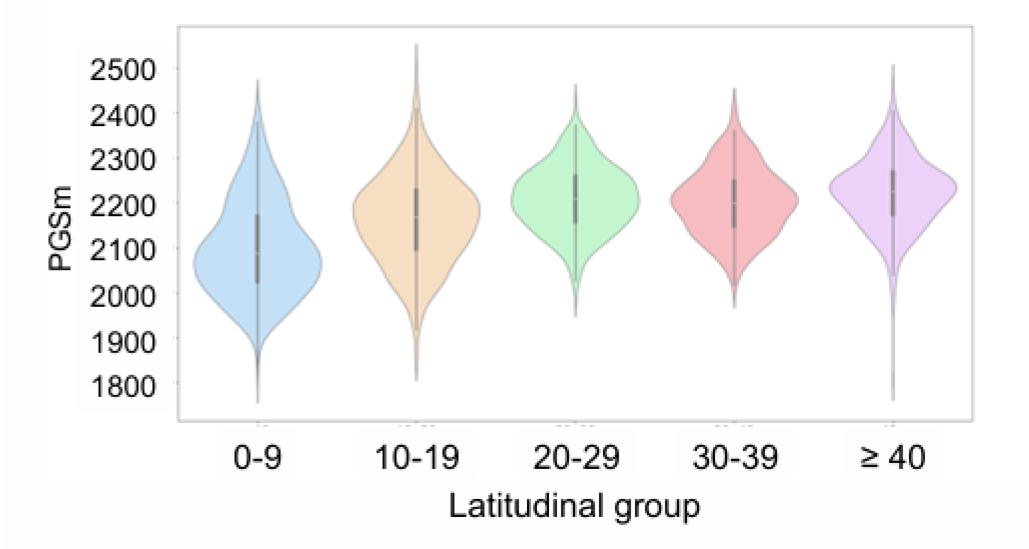


Supplementary Fig. 2. Distributions of PGSm in latitudinal groups, including the populations with migrations into low latitudes. Populations belonging to each latitudinal group are as follows: LWK, CLM, STU, YRI, MSL, ESN, and KHV for 0°–9°; PEL, ACB, GWD, ITU, and PUR for 10°–19°; CDX, MXL, and CHS for 20°–29°; GIH, BEB, PJL, and JPT for 30°–39°; CHB, IBS, TSI, GBR, and FIN for ≥ 40°. The Bonferroni-corrected threshold is $P$ = 0.0125 for the four Kruskal-Wallis tests (see also Fig. 2). Regarding the PGSm distributions, a significant difference in the trend towards morning chronotype was detected among the latitude groups. The PGSm distribution shifted higher in the three latitudinal groups above 20°N than in two groups below 20°N ($P = 3.4 \times 10^{-100}$; Kruskal-Wallis test for all latitudinal groups). Furthermore, the trend towards the morning chronotype was more pronounced in the ≥ 40° latitudinal group ($P = 1.2 \times 10^{-3}$; Kruskal-Wallis test among the 20°–29°, 30°–39°, and ≥ 40° latitudinal groups).

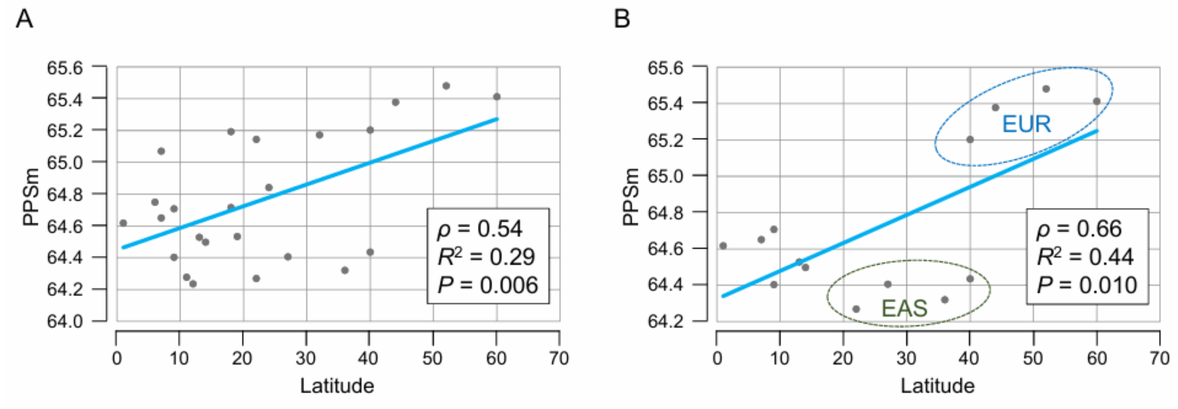


Supplementary Fig. 3. Association between latitude and frequencies of positive-effect alleles of HEI. (A) PPSm distributions. (B) PPSm distribution after the exclusion of populations with migrations into low latitudes. EUR, populations from Europe; EAS, populations from East and Southeast Asia.

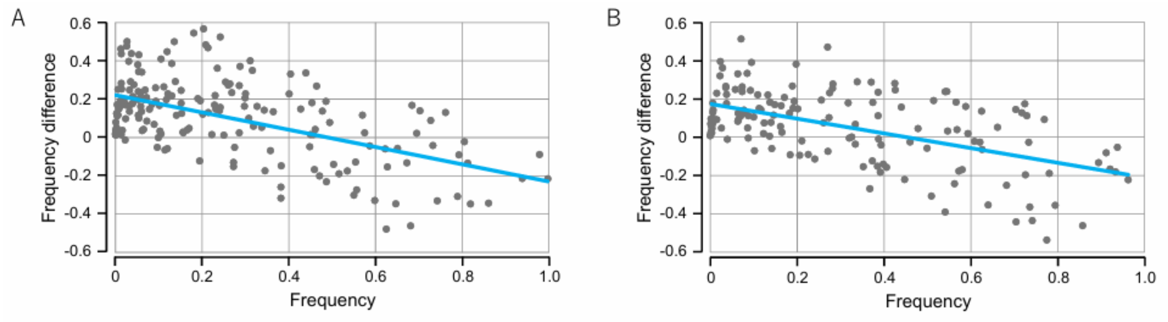


Supplementary Fig. 4. Allele frequency differences between populations living at < 20°N and > 20°N. (A) The morningness alleles. (B) The eveningness alleles.

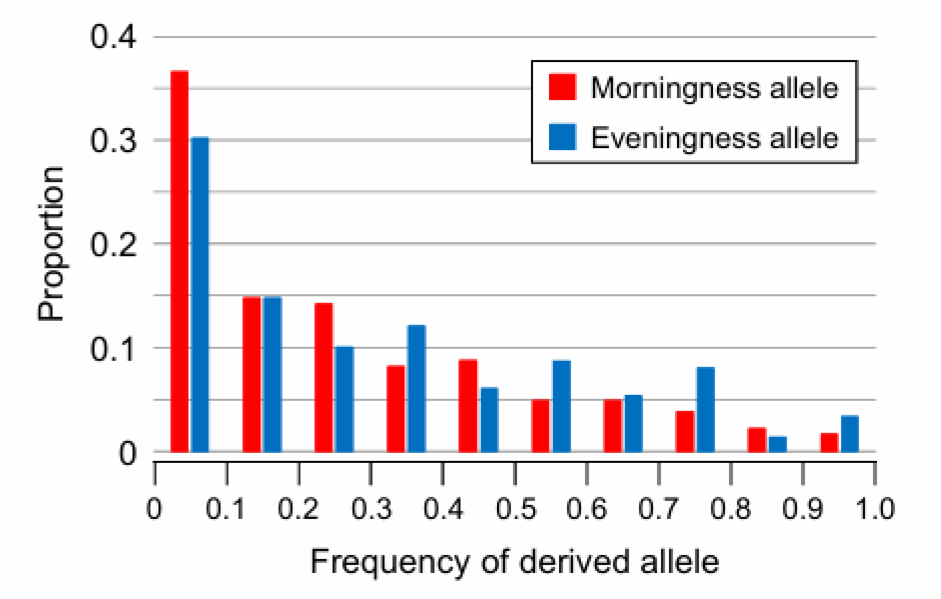


Supplementary Fig. 5. Derived allele frequencies in Africa.

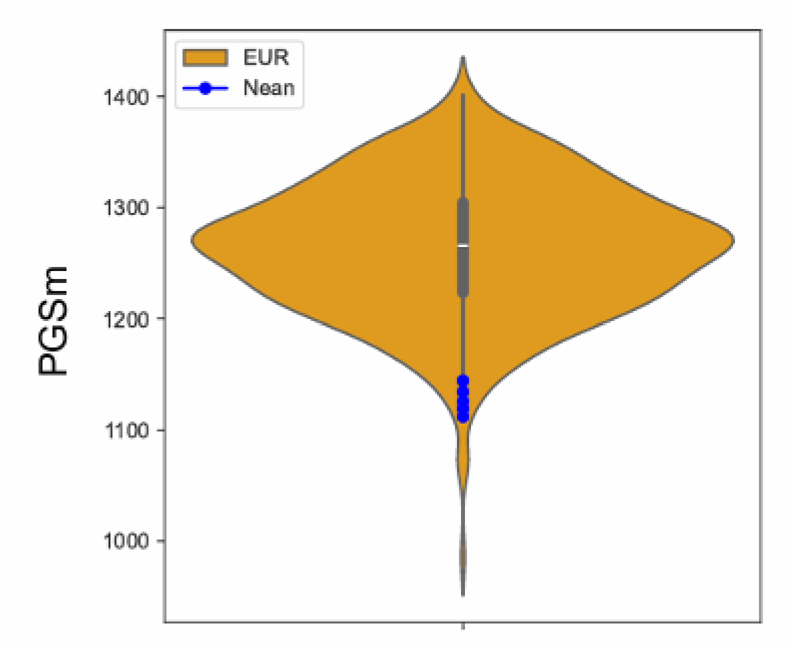


Supplementary Fig. 6. PGS distributions. PGSs of Europe in the pdMHs are standardised and shown by violin plots. PGS values of Neanderthal individuals are represented by blue circles.

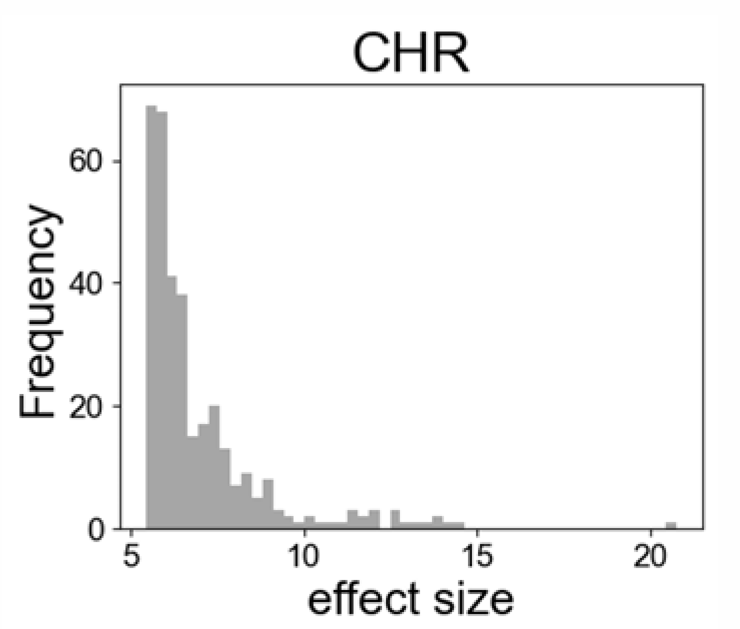


Supplementary Fig. 7. Distribution of effect sizes of positive-effect alleles of CHR.

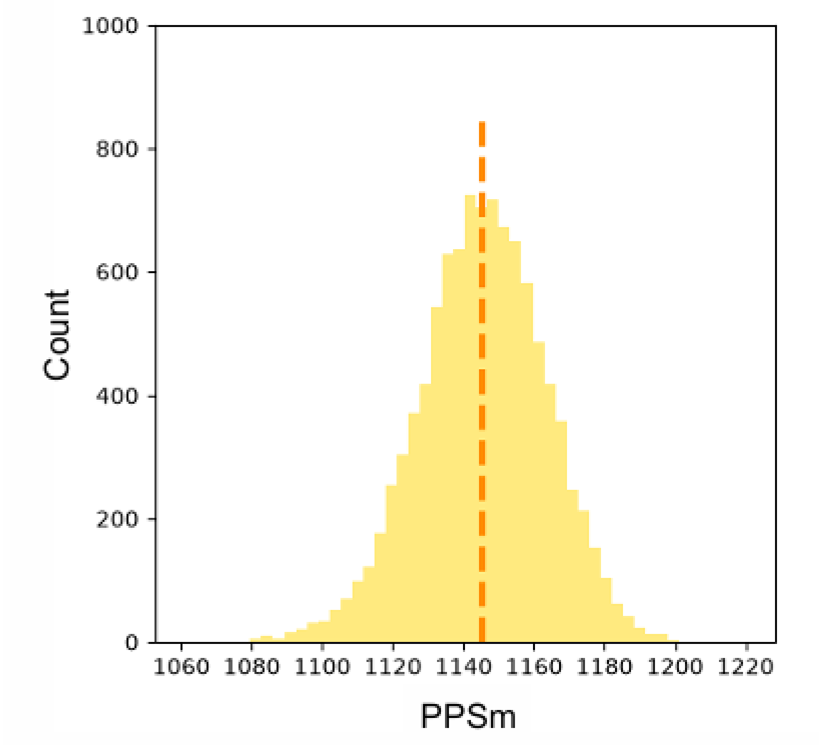


Supplementary Fig. 8. Distribution of PPS values estimated using five individuals randomly selected from EUR and actual PPS value of EUR (vertical dotted line). PPS distribution was generated by 10,000 replications.

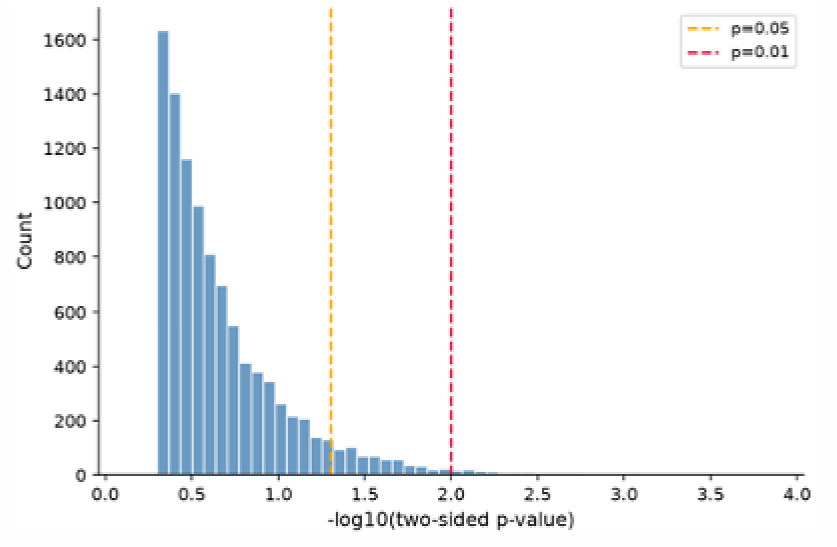


Supplementary Fig. 9. Comparison between actual PPS and the inferred PPS distribution. We selected a number of alleles from Europe for each SNP that was equal to the number of alleles available from five Neanderthal individuals, and obtained a possible distribution inferred from the frequencies of morningness alleles in the selected alleles. The comparisons were replicated 10,000 times.

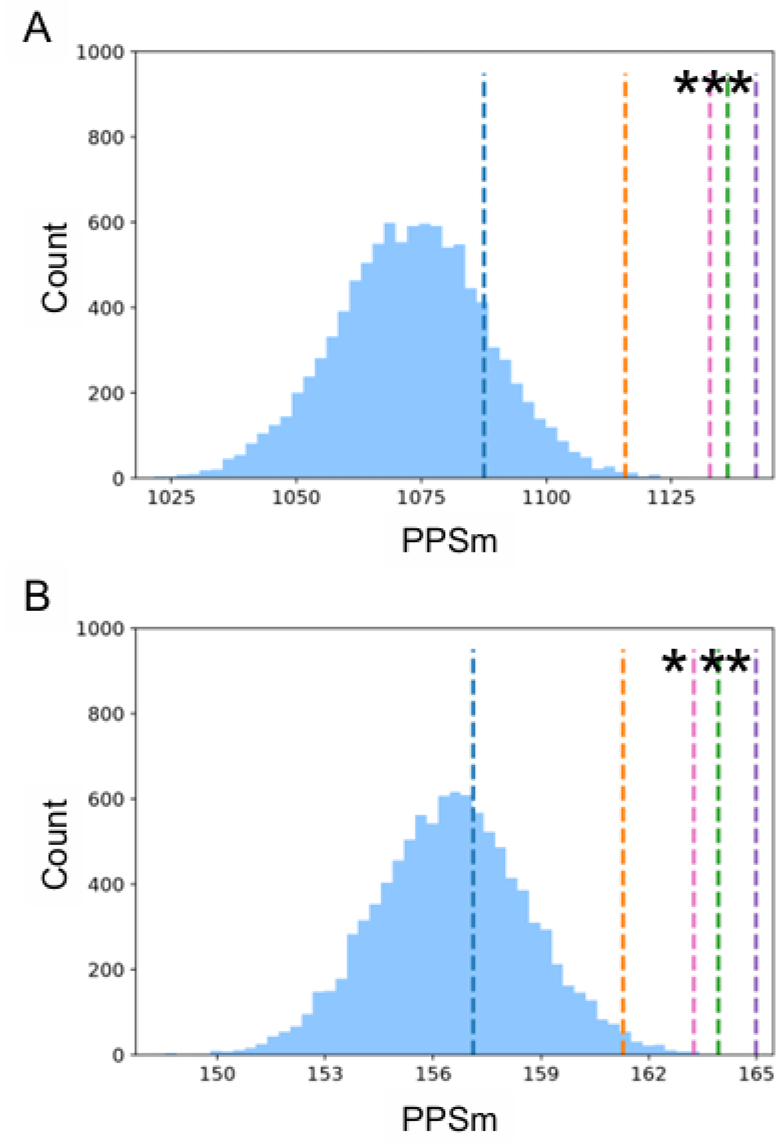


Supplementary Fig. 10. Comparison of chronotype trends between AMHs and Neanderthals. (A, B) PPSm values. PPSm values of the latitudinal groups (0°–9°, 10°–19°, 20°–29°, 30°–39°, and ≥ 40°) were estimated, and are shown as vertical dotted lines in blue, orange, green, pink, and purple, respectively. The PPSm for Neanderthals is shown by a possible distribution. PPSm estimation was performed with effect sizes from GWAS (A) and with homogeneous effect sizes (B). Asterisks indicate a Bonferroni-corrected significant difference compared with the Neanderthal distribution.
